# Thought Control Failure: Sensory Determinants and Functional Effects

**DOI:** 10.1101/293761

**Authors:** Eugene Lee Kwok, Gaelle Leys, Roger Koenig-Robert, Joel Pearson

## Abstract

The ability to control one’s thoughts is important for mental wellbeing, attention, focus, future planning and ideation. While there is a long history of research into thought control, the inherent subjectivity of thoughts has made objective investigation, and thus mechanistic understanding difficult. Here, we report a novel method to empirically investigate thought control success and failure by objectively measuring the sensory strength of visual thoughts. We use the perceptual illusion binocular rivalry to assess emergent images in mind during two common thought control strategies: thought suppression and thought substitution. Thought suppression was ineffective, suppressed thoughts primed subsequent rivalry dominance at the same level as intentionally imagined thoughts. This priming effect was disrupted by concurrent uniform luminance and changes in retinotopic location, suggesting early visual representations/traces. While individuals showed some metacognition of thought suppression, strikingly, the perceptual effects remained even when thoughts were reported as successfully suppressed, indicating these thoughts may exist outside of reportable awareness. In contrast, thought substitution was more effective in controlling the perceptual effects and showed good metacognition. A thought control index predicted greater levels of trait mindfulness, while high levels of anxiety and schizotypy were related to poor thought control. Overall, our findings offer a novel method to track thoughts before and after they emerge into awareness and suggest that non-reportable and involuntary thoughts form visual representations pivotal to thought control failure.

## Significance Box

Unlike bodily motor control, controlling one’s thoughts, despite its importance for multiple dimensions of everyday life, has remained largely enigmatic. The inherent subjectivity of our thoughts has made objective examination difficult. In this study we report a novel method using the visual illusion binocular rivalry to measure the sensory trace of thought control and its failure. We find evidence of a sensory thought trace even when individuals report having successfully suppressed a thought. While the technique of thought substitution did manage to eliminate the sensory trace. Our study provides a unique method to objectively investigate the mechanisms of thought control and its failure across different populations and contexts.

The ability to control our own thoughts, is an important mental capacity and is important for attention, focus, future planning and ideation [1-4]. Given that up to 80% of the general population experience some form of unwanted intrusive thoughts [5], the ability to control thoughts is also an important determinant of mental wellbeing [6]. Indeed, failure to control thoughts has been linked to various mental disorders including anxiety [7, 8], obsessive-compulsive disorder (OCD) [9, 10] and schizophrenia [11, 12].

Since the work of Freud, the idea of voluntarily repressing a thought, thought suppression, has become an active phrase in the general populace and has attracted much attention, despite the lack of any clear objective methodology to investigate the phenomenon. Attempting to control thoughts by direct thought suppression (*not* thinking about a given thought) is often difficult and subjective reports suggest suppression mostly fails [6]. Instead of directly eliminating thoughts from the mind, reports suggest suppression paradoxically leads to heightened preoccupations with these thoughts [13]. For example, individuals instructed to suppress the thought of a white bear are often unable to do so and report intrusions of the thought inadvertently arising a short time later [14]. Research into thought suppression failure (e.g. [15-20]) and its consequences have been studied in a wide variety of domains including organisational behaviour [21], smoking addiction [22], immune function [23] and psychopathology [24-27]. However, largely due to the inherent private subjectivity of thought suppression, its existence as a possible process, dynamics, where and why such suppression failures originate, remains largely unknown.

While thought suppression is the most commonly studied thought control strategy, other forms of thought control have been examined and seem to be more effective. For example, thought substitution, in which a suppressed thought is instead replaced by a substitute thought, has shown evidence for reduced thought intrusion frequency and thought control failure [6, 14, 28]. More recently, mindfulness has shown evidence as an effective form of mental control [29]. While these forms of thought control seem more effective than direct thought suppression, without a mechanistic understanding of these processes, research into why one method might work over another, and indeed if there are potentially more effective methods remains limited.

Wegner [13] first attributed the inability to control thoughts using thought suppression to the interaction between two practical, but simultaneously conflicting, mental processes. The first involves a conscious process that attempts to achieve a state of mind free from the to-be-suppressed thought. The second is an unconscious monitoring process that checks for unwanted instances of the to-be-suppressed thought. Wegner proposed thoughts unwillingly enter consciousness due to the ironic opposition between these two processes, ultimately leading to thought control failure. Neuroimaging evidence has generally supported this view [30-32]. However, while mechanisms have been proposed, the empirical investigation and objective measurement of thought control has been limited.

The inherent subjectivity of thoughts has further made the objective examination of thought control difficult. For the most part, previous research has used subjective self-reports to examine thought control (e.g. [14, 28, 33, 34]). While self-report measures have provided valuable insight into the examination of thought control, these measures may be prone to bias, experimenter demand or social desirability effects [35], and thus make it difficult to examine the underlying mechanisms.

To overcome this subjectivity and to shed light on the underlying mechanisms of thought control and its failure we devised a novel method to objectively study the control of visual thoughts using the illusion binocular rivalry. Binocular rivalry is a visual illusion that arises when two different images are presented one to each eye, inducing bistable perceptual alternations between the two images [36]. Binocular rivalry has been used to objectively measure the sensory strength of mental images [37-40] and prior perceptual stimuli [37, 39, 41]. Here we devised a novel method to utilise a brief binocular rivalry presentation to objectively assess visual representations that might emerge during attempts at thought control.

In each trial, we instructed participants to either imagine, suppress or substitute the thought of a red or green coloured object, before being presented with a brief red-green binocular rivalry stimulus. Any priming for imagined, suppressed or substituted thoughts could then be calculated to provide an objective measure of the sensory strength of these visual thought representations. To probe subjective metacognition of thought control, we instructed participants to report when thought control failed and compared these subjective reports to their level of rivalry priming. Lastly, a Thought Control Index was devised to investigate the relationship between thought control and four psychological traits: anxiety, obsessive-compulsive tendencies, schizotypy and mindfulness.

Results showed suppressed thoughts led to binocular rivalry priming significantly above chance (50%) and almost as strong as priming for intentionally imagined thoughts, indicating a failure to control the sensory trace of thoughts using thought suppression. Surprisingly, priming for trials reported as successfully suppressed was still significantly above chance, suggesting the possibility of emergent non-conscious sensory representations during attempts at thought control. Thought substitution was more, although not completely effective, in controlling the sensory trace of visual thoughts. Individuals with high trait mindfulness exhibited greater levels of thought control, while those with high anxiety or schizotypy traits exhibited lower levels of thought control. Interestingly, there was no relationship with obsessive-compulsive tendencies. Our data suggest that thought control failure is linked to the formation of sensory representations during attempts at thought control.

## Methods

Ten participants (3 female, M = 25.60 years, SD = 4.62), completed the thought suppression, luminance, location and thought substitution experiments. Ten participants were chosen to reflect the number of participants in past research which used binocular rivalry to measure the sensory strength of visual thoughts [37, 39, 42]. All participants were the same across these four experiments. However, two participants who were unable to complete the location experiment were replaced by two age and gender matched participants. The psychological traits experiment consisted of a total of 67 participants (35 female, M = 21.88, SD = 5.03), which included the data from the ten participants in the main experiment. 67 participants were chosen to reflect the number of participants used in past correlational research on individual differences in the strength of mental imagery (visual thoughts) [43].

### Thought suppression experiment

All experiments followed the design of the thought suppression experiment (*Figure 1A*). The task was carried out on a Windows 7 PC running Psychophysics Toolbox 3 [44] in MATLAB on a 85Hz Dell Trinitron P1130 CRT monitor 1280×1024 resolution. Red-green coloured filters, one over each eye, were worn by participants to view the binocular rivalry stimulus – a red-green Gabor patch. A chinrest positioned the participant’s head 57cm from the screen. Before the experiment, each participant completed an eye dominance test (as reported in [37, 42]) to determine the stimulus intensity at which the red and green grating stimuli were equally dominant for each participant.

**Figure 1.**
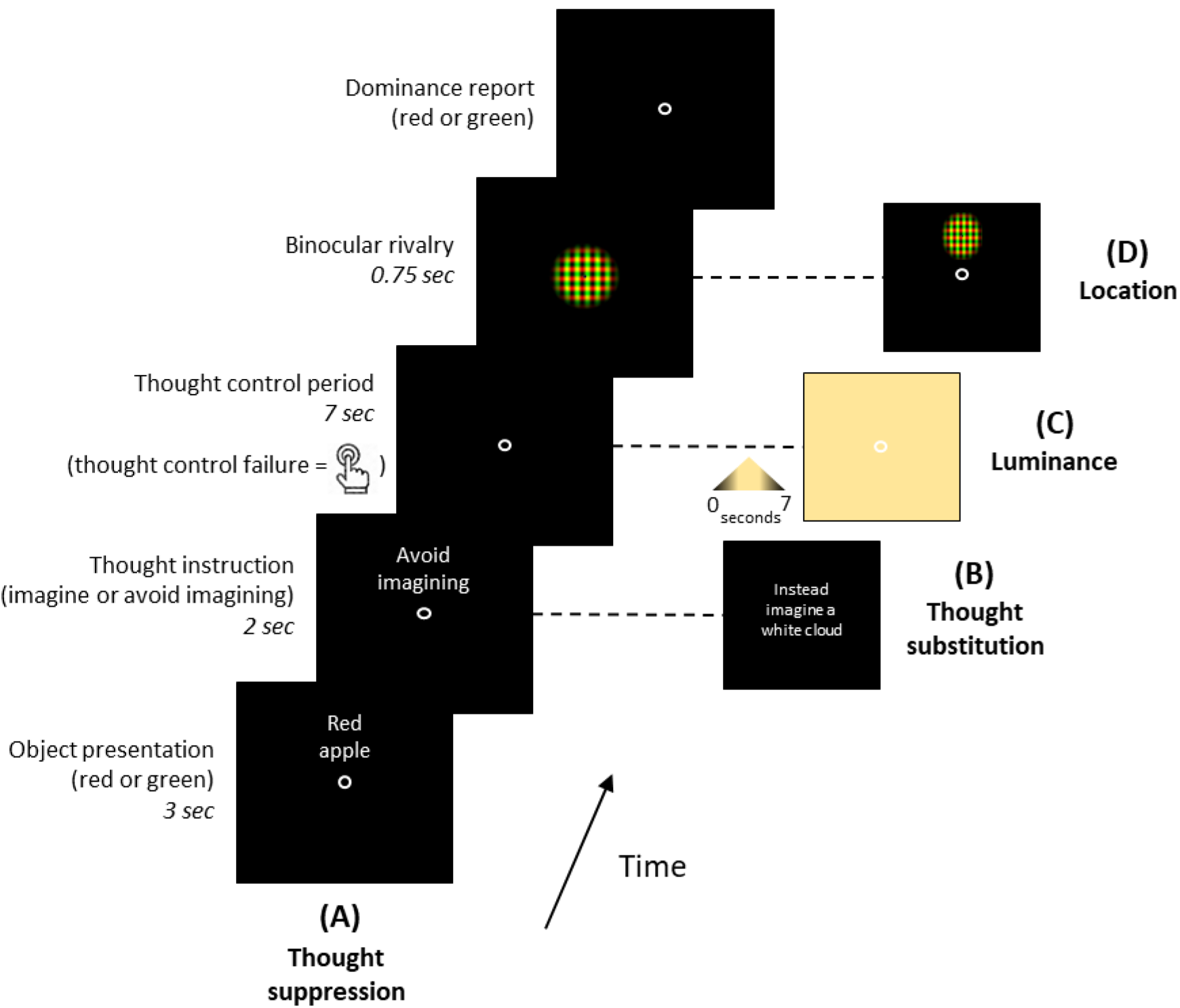
Measuring thought control with binocular rivalry– time course and conditions. **(A)** In the thought suppression experiment, participants were presented with one of three red (red apple, red chilli, red tomato: shown prior to the experiment) or green (green broccoli, green cucumber, green lime) written object cues and were then instructed to either ‘imagine’ or suppress (‘avoid imagining’) the thought of the object for 7 seconds. In suppression trials, participants pressed the assigned button to indicate occurrence of the thought– a failed suppression attempt – during the suppression period. Participants were then briefly presented with a red-green rivalry stimulus and reported dominance. **(B)** In the thought substitution experiment, participants instead imagined a white object (white cloud, white sheep, white cauliflower). **(C)** In the luminance experiment, participants viewed changing background luminance during the imagery/suppression period. **(D)** In the location experiment, participants viewed the rivalry stimulus in one of four locations (top, bottom, left, right) around the screen (only top location shown). Two additional questions on vividness and effort were presented at the end of each trial and were rated on a scale of 1-4. Note: vividness, effort, memory control and decision bias control questions not shown.

The experiment consisted of a minimum of three blocks of 42 trials (minimum total of 126 trials) over a one-hour session. If time permitted, additional blocks were performed. Participants focused on a fixation point at the centre of the screen for each trial and a black background was present throughout the experiment. Each trial comprised of four stages: (1) object presentation (2) thought instruction (3) thought control period (4) binocular rivalry.

In the object presentation stage, the words of one of six object stimuli (three green objects – green broccoli, green lime, green cucumber; three red objects – red chilli, red apple, red tomato) were pseudo-randomly presented on the screen as a cue. Next, in the thought instruction stage, the words ‘imagine’ or ‘avoid imagining’ appeared on the screen. In the ‘imagine’ condition, participants actively imagined the presented object. In the suppression or ‘avoid imagining’ condition, participants attempted suppression of any thoughts of that object. The presentation order of the two conditions was randomised to remove order effects and the two conditions were intermixed rather than presented in separate blocks to increase participant engagement in the task. Then, in the thought control period, participants either imagined or suppressed the thought of the object stimulus for seven seconds, depending on the thought instruction provided in the previous stage. Importantly, if participants failed at thought suppression they were instructed to press the spacebar immediately. Lastly, in the binocular rivalry stage, a red-green Gabor patch appeared (red horizontal: luminance = 13.2 cdm^2^, CIE *x* = 0.360, *y* = 0.358; green vertical: luminance = 10.9 cdm^2^, CIE *x* = 0.263, *y* = 0.568) and participants perceived binocular rivalry between the colours red and green (.75s). Participants indicated, by pressing assigned keys, whether they perceived colour dominance in (a) red (b) green (c) ‘mixed’ (50% red/green).

Each block of 42 trials contained 36 experimental trials. Within the 36 experimental trials, there were an equal number of imagery and suppression trials (18 ‘imagine’, 18 ‘avoid imagining’) and trials of the object stimuli were divided equally amongst the six objects (6 ‘red apple’, 6 ‘green cucumber’ etc. – total of 18 red objects, 18 green objects). The remaining 6 trials consisted of memory control trials. These trials required participants to correctly identify the cue object presented to them during the object presentation stage i.e. the object they were instructed to imagine or suppress, from a list of three object stimuli. These questions encouraged participants to pay attention to the stimuli to be imagined/supressed. This control ensured participants were not ignoring the instructions in the suppression condition. The order in which the 42 trials were presented was randomised to eliminate order effects.

### Luminance experiment

In the luminance experiment (*Figure 1C*), we examined whether the sensory trace of suppressed thoughts could be disrupted by bright uniform background luminance. The procedure for the luminance experiment was identical to the main thought suppression experiment except for one modification. During the imagery/suppression period, the background luminance smoothly ramped up from black to yellow (mix of the red and green) background (luminance = 10.8 cdm^2^, CIE *x* = 0.375, *y* = 0.546) over two seconds, remained a yellow background for three seconds, and then ramped back down to a black background (total of seven seconds). Yellow was chosen as the background luminance colour as it was a mix between the red and green colour of the stimuli and thus would not result in any bias during binocular rivalry.

### Location experiment

In the location experiment (*Figure 1D*), we examined whether changing the location of the binocular rivalry stimulus would affect any priming we observed from suppressed thoughts. The procedure for the location experiment was identical to the main thought suppression experiment except for three modifications. (1) The binocular rivalry stimulus was presented at one of four randomised locations from fixation (top, bottom, left, right) instead of at fixation. (2) A random 20% of trials were decisional bias control trials, in which a physically fused and equally dominant red-green Gabor patch was presented on the screen in place of the standard binocular rivalry stimulus, to measure any potential decisional bias (as reported in [37]). These trials monitored whether participants were accurately reporting binocular rivalry dominance (3). Two additional questions were included at the end of each trial. A vividness question asked participants to rate how vivid the thought of the object was during the imagery/suppression period. The vividness question appeared in all imagery trials but was only asked in failed suppression trials i.e. when participants reported thought suppression failure with a button press. Participants were also asked to rate the amount of effort put into imagining or suppressing the thought. Vividness and effort were rated on a scale of 1-4 (1 = not vivid/effortful, 4 = very vivid/effortful).

### Thought substitution experiment

In the thought substitution experiment (*Figure 1B*), we examined the effects of thinking about a substitute object stimulus, instead of the original object stimuli, on rivalry priming. The procedure for the thought substitution experiment was identical to the main experiment except for one modification. In the thought instruction stage, participants were presented with the instruction to imagine a white object (either a white cloud, white cauliflower or white sheep) instead of the instruction ‘avoid imagining’.

### Psychological traits experiment

The psychological traits experiment followed the same procedure as the main thought suppression experiment (*Figure 1A*). Four traits – anxiety, obsessive-compulsive tendencies, schizotypy and mindfulness – were examined. Individual dispositions were measured through four standard questionnaires: the second half of the State Trait Anxiety Inventory (STAI) (measuring only trait anxiety and not state anxiety, which was not relevant for this study) [45], the Obsessive-Compulsive Inventory–Revised (OCI-R) [46], the Oxford-Liverpool Inventory of Feelings and Experiences (O-LIFE) [47] and the Mindfulness Attention Awareness Scale (MAAS) [29].

### Statistical analysis

Participants with poor performance (chance or below) on the memory control trials and decision bias control trials (in the location experiment) were removed from the analysis as they could not be trusted to have been attending to the task or accurately reporting binocular rivalry dominance. This amounted to seven participants being removed in the psychological traits experiment due to poor performance on the memory control trials. All participant performed well on the decision bias control trials and thus no participants were removed from the analysis in the location experiment.

Four participants with priming levels below 40% for imagined or suppressed thoughts in the psychological traits experiment were also removed because we were unable to determine the sensory strength of their thought representations. In *Figure* 2A, priming for each participant was measured by calculating the number of trials in which the dominant rivalry colour matched the colour of the imagined, suppressed or substituted stimulus, and then dividing by the total number of imagery, suppression or substitution trials respectively. When calculating priming for successful and unsuccessful suppression trials in *Figure 2B*, the distribution of results did not allow the use of standard statistics as the number trials with button presses varied across individuals. Thus, all the trials for each participant were pooled and resampling statistics (showing 95% confidence intervals) were conducted to calculate priming. In all analyses, family-wise error in multiple comparisons were controlled by setting the false discovery rate (FDR) at *q*=0.05 [48].

**Figure 2.**
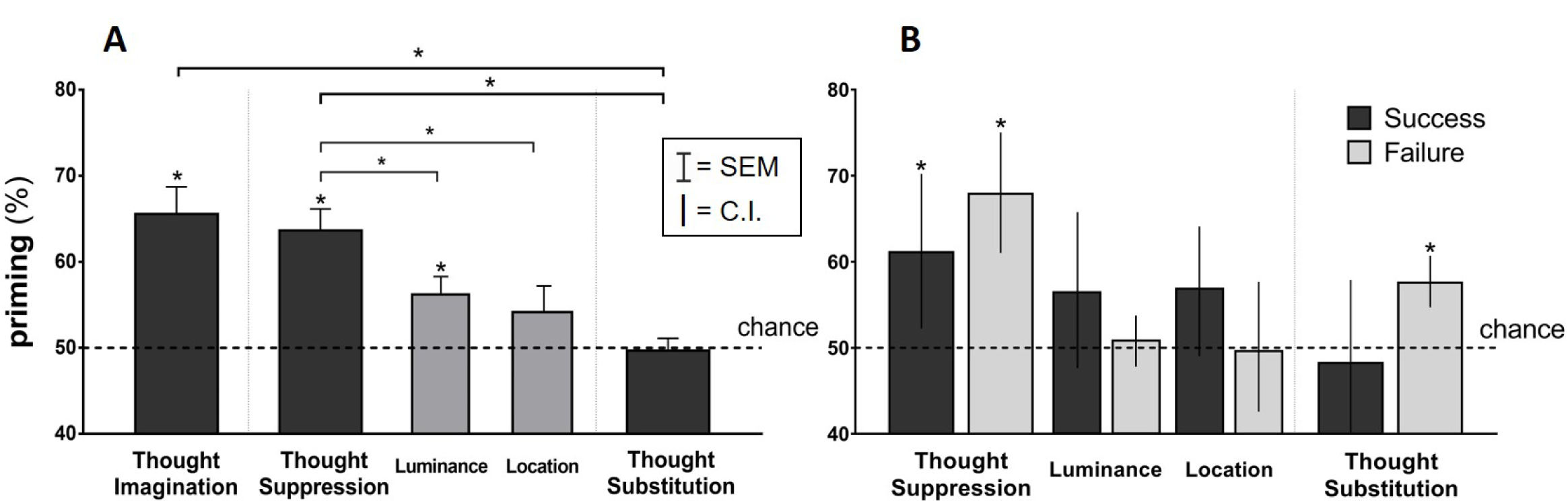
(A) Sensory strength of imagined, suppressed and substituted thoughts. In the thought suppression experiment both the suppression and active imagery conditions showed significant priming. There was no significant difference in between imagined or suppressed conditions. In the luminance and location experiments, priming was significantly reduced compared to the thought suppression experiment suggesting suppressed visual thoughts were represented in visual areas. The thought substitution strategy experiment was effective in reducing priming to chance levels. Error bars represent standard error of the mean (+SEM). **(B) Metacognition of thought control success and failure**. Sensory priming varied as a function of self-reported thought suppression success or failure. Error bars represent 95% confidence intervals. Multiple comparisons were corrected by controlling the false discovery rate (FDR) at *q*=0.05.

## Results

### Thought Suppression

We examined whether suppressed thoughts formed a quantifiable sensory trace by measuring how they might affect subsequent rivalry presentations. We found above chance binocular rivalry priming for suppressed thoughts (t(9)=5.893, p<0.001), with no difference in priming compared to actively imagined thoughts (p>0.05), which also exhibited above chance priming (t(9)=5.176, p<0.001) (*Figure 2A*). In other words, when participants attempted to not think about a red object, they were significantly more likely to report red in the subsequent rivalry presentation. This indicates, perhaps surprisingly, that thought suppression was ineffective at preventing the sensory trace of visual thoughts from forming. The probability of self-reported thought suppression failure, as indicated by participant button presses, during the main thought suppression experiment was 0.29 and was significantly above zero (t(9)=2.80, p=0.0433) (*Supplementary Figure 1*).

To test if the priming during attempted thought suppression was of a sensory nature we utilised uniform background luminance as a potential perturbation to the formation of a sensory representation. Luminance is a visual feature strongly represented in early visual areas [49, 50], including the primary visual cortex (V1) [51], a uniform luminance background can disrupt the formation of intentional mental images [37, 38, 42]. Luminance disruption would thus be expected to result in reduced priming, if suppressed visual thoughts are represented in visual areas. Priming for suppressed thoughts in the luminance condition were significantly lower compared to priming with a black background (t(18)=2.231, p=0.041), but still above chance (t(9)=2.725, p=0.025) (*Figure 2A*), suggesting that priming from suppressed thoughts most probably involves sensory representations in visual cortex. The probability of self-reported thought suppression failure in the luminance experiment was 0.147 and was significantly above zero (t(9)=2.387, p=0.0444), but lower compared to the main experiment, although not significant (t(18)=1.188, p=0.2154) (*Supplementary Figure 1*).

Next, we further examined the sensory nature of the priming effect from suppressed thoughts by investigating the local retinotopic nature of the priming bias. If the priming effect was local in visual space, this would suggest that suppressed thought representations are represented in retinotopically organized visual areas. We tested this by changing the spatial location of the binocular rivalry presentations. Spatial location is a visual feature processed retinotopically [52-54], and this retinotopic organisation is preserved during mental imagery [55, 56]. Consequently, probing the effects of thought suppression at a different retinotopic location (by changing the location of binocular rivalry) would be expected to decrease priming [37, 39]. After changing the location of binocular rivalry, priming for suppressed thoughts was significantly lower compared to priming in the main thought suppression experiment (t(18)=2.727, p=0.022), and was not significantly different from chance (p>0.05) (*Figure 2A*). The probability of self-reported thought suppression failure was 0.46, which was significantly above chance (t(9)=4.757, p=0.0059) and comparable to that of the main thought suppression experiment (p>0.05) (*Supplementary Figure 1*). Combined with the findings from the luminance disruption experiment, these results provide evidence that attempted suppressed thoughts are likely represented in visual areas, including early visual cortex.

In the location experiment, we also explored participant ratings of vividness and the amount of effort used when imagining or suppressing a thought, both rated on a scale of 1-4. As shown in *Supplementary Figure 2*, vividness of imagined thoughts was significantly stronger compared to the vividness of suppressed thoughts that arose during failed suppression trials (t(16)=2.998, p=0.009). No difference was found in the amount of effort expended in imagining or suppressing a thought (p>0.05), and there was no significant difference in effort between successful and failed suppression trials (p>0.05; *Supplementary Figure 2*). The location experiment additionally contained decision bias control trials, which all participants answered correctly (mean of >90% correct), indicating accurate reporting of binocular rivalry dominance.

To examine metacognition of thought suppression, we compared the subjective reports of thought suppression success or failure with the level of sensory priming. Priming was greater for trials in which thought suppression was reported as a failure compared to trials reported successful, although this relationship was not significant (p=0.18), suggesting a trend towards some metacognition. However, priming was significantly above chance for both successful and failed suppression trials (p=0.014, p=0.001, resampling statistics; *Figure 2B*), indicating the sensory representations of successfully suppressed thoughts were strong enough to drive sensory priming despite the absence of reported awareness. As expected, there was no difference in priming between successful and failed thought suppression trials in the luminance and location experiments (p>0.05) and neither were significantly different from chance (p>0.05) (*Figure 2B*).

### Thought Substitution

We investigated if imagining a substitute thought instead of the object to be suppressed would be an effective strategy for reducing the perceptual influence of visual thoughts. Previous research has shown that a focused distraction strategy, in which individuals focus on a distractor or substitute thought instead of a target thought, can reduce the frequency of reported thought control failures [14, 57]. If this is the case, we might expect priming for substituted thoughts to be lower than that of imagined and suppressed thoughts. Participants were instructed to imagine white objects as a replacement strategy, to prevent thoughts about the coloured objects emerging (see methods).

Priming in the thought substitution condition was significantly lower than the standard thought suppression condition (t(9)=3.995, p<0.001) and imagined thoughts (t(9)=4.537, p<0.001), and was not significantly above chance (p>0.05) (*Figure* 2A), suggesting thought substitution was effective in reducing the sensory trace of visual thoughts. However, failed thought substitution trials (in which the coloured object thought still emerged) showed above chance priming (p<0.001, resampling statistics; *Figure 2B*) compared to successful thought substitution trials which showed no priming effect (p>0.05). This suggests thought substitution was ineffective when the strategy failed, but that participants had good metacognition of when such failure occurred. The probability of thought substitution failure was 0.132 and was significantly above chance (t(9)= 2.365, p=0.0444).

### Thought Control Index

We devised a Thought Control Index to measure individual ability to control thoughts using thought suppression. Scores on this index were calculated by finding the difference in priming between imagined thoughts (imagery priming) and suppressed thoughts (suppression priming) 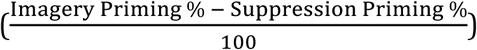 for each participant. Positive index scores indicate the sensory strength of imagined thoughts was stronger than suppressed thoughts, suggesting the ability to control visual thoughts. Zero or negative index scores indicate the sensory strength of suppressed thoughts was as strong or greater than that of imagined thoughts, suggesting the inability to control visual thoughts. We did not simply use suppression priming as a measure of thought control because this measure would not take into account individual differences in mental imagery strength. Since thought suppression priming was strongly correlated with imagery priming (r=0.52, p<0.001; *Supplementary Figure 4*), this suggested that a good individual measure of thought control might be the degree to which suppression priming differed relative to imagery priming.

To confirm that Thought Control Index scores were assessing the relative effectiveness of thought control, we applied it to the findings in the thought suppression, luminance, location and thought substitution experiments. As expected, index scores during thought substitution were significantly higher than index scores during thought suppression (t(9)=4.208, p=0.002), and were significantly above zero (t(9)=3.929, p=0.004) (*Supplementary Figure 3*), and thus confirmed the validity of the index.

**Figure 3.**
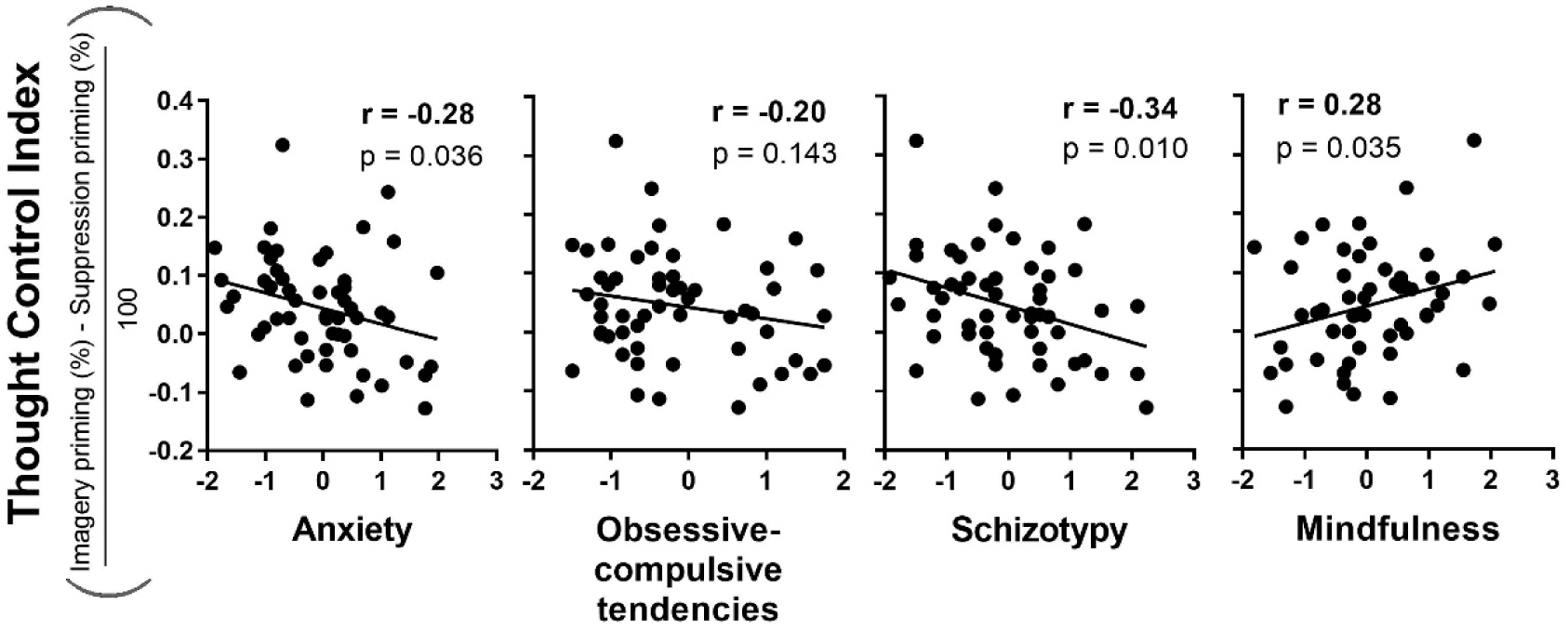
Thought Control Index and psychological traits (n=56). Thought Control Index scores were correlated with three psychological traits: anxiety, schizotypy and mindfulness, but not obsessive-compulsive tendencies.

Next, we used the index to explore individual differences in the ability to control visual thoughts across four psychological traits (anxiety, obsessive-compulsive tendencies, schizotypy and mindfulness; see methods for questionnaire information) (*Figure 3*). For 56 participants (comprising data from 46 new participants and data from the 10 participants in the main thought suppression experiment), Pearson correlations with index scores revealed negative correlations with anxiety (r=-0.28, p=0.036) and schizotypy (r=-0.34, p=0.010) indicating high levels of trait anxiety and schizotypy predicted lower levels of thought control. A positive correlation was found with mindfulness (r=0.28, p = 0.035), indicating high levels of mindfulness predicted greater levels of thought control. No significant correlation was found between index scores and obsessive-compulsive tendencies (r=-0.20, p>0.05).

Further analyses showed that index scores across all participants followed a normal distribution (min: −0.127, max: 0.324, SD = 0.09; *Supplementary Figure 5*). The distribution was centred around a mean of 0.045, which was not significantly above zero (p>0.05), indicating participants were, on average, poor at controlling the sensory influence of suppressed visual thoughts.

We verified the validity of our questionnaire data with a correlation matrix for the four traits (*Supplementary Figure 6*), which confirmed strong intercorrelation between the ‘negative’ traits (anxiety, obsessive-compulsive tendencies, schizotypy) as previously reported [58-60], and additionally, that these negative traits were inversely correlated with the ‘positive’ trait of mindfulness, which has also been previously reported [61-63].

### Memory control trials

Memory control trials were present in all experiments and monitored if participants were attending to the object stimuli cues. Aside from seven participants in the Psychological Traits experiment who performed poorly (below chance levels) and were removed from the analyses, all participants showed good performance on the memory control trials (mean of >85% correct).

## Discussion

Here, we utilised the visual illusion binocular rivalry to measure the sensory trace of emergent thoughts during attempts to control and supress them. Binocular rivalry priming for suppressed thoughts was significantly above chance and was, surprisingly, comparable to priming for actively imagined thoughts (*Figure 2A*). This suggests the sensory representations of suppressed visual thoughts were as strong as those of imagined thoughts. This bias from supressed thoughts showed the sensory characteristics of low-level visual areas, being disrupted by changes in luminance and the spatial location of the rivalry stimulus. Strikingly, even when participants reported successful thought suppression, subsequent rivalry was still significantly biased, suggesting the perceptual effects of emergent sensory representations were adequate to bias rivalry, but not trigger a subjective report of suppression failure. This is important because trials in which suppression was believed to be successful comprised nearly 70% of the total number of suppression trials (*Supplementary Figure 1*), indicating for the majority of trials, individuals believed they were successful in thought suppression. The apparent ease in which individuals felt they could suppress thoughts was reflected in the subjective ratings of effort and vividness. There was no difference in the amount of effort expended between imagined and suppressed thoughts and during suppression failure, the vividness of suppressed thoughts was rated as less vivid than imagined thoughts (*Supplementary Figure 2*).

It is important to recognise that the data suggests that the priming effects we observed for suppressed thoughts were sensory in nature, and not simply due to high-level decisional bias or semantic priming. In the luminance and location experiments, priming was significantly *below* that in the main thought suppression experiment (*Figure 2A*). Since luminance and location are visual features represented mainly in low-level visual areas [49, 50], including the primary visual cortex (V1) [51], the observed perturbations suggest the ‘suppressed’ thought representations were sensory in nature (i.e. suppressed visual thoughts have perceptual representations). This suggests the priming effect was driven by the activation of sensory areas which subsequently primed rivalry dominance.

Thought substitution, in which participants substituted the thought of a red or green object with a white object, was more effective in controlling visual thoughts compared to thought suppression. Priming for substituted thoughts was significantly lower than both imagined and suppressed thoughts, and was not significantly different from chance (*Figure 2A*). This suggests thought substitution may work to neutralize the emergent sensory trace of the unwanted thought. However, while these results reflect the effectiveness of employing a thought substitution strategy to improve thought control [14], thought substitution was not completely effective. When thought substitution attempts failed (i.e. when individuals failed to substitute the white object stimuli), the priming effect returned (*Figure 2B*), indicating the strategy was only effective during successful substitution. However, individuals showed good metacognition of when their thought substitution attempts were successful or failed. While more effective than thought suppression, thought substitution failed on average 13% of the time (*Supplementary Figure 1*), thus, thought substitution did not completely eliminate the probability of thought control failure. Considering both the thought suppression and thought substitution experiments, these results suggest that neither thought control strategy was completely effective in controlling visual thoughts.

Our data suggests that when individuals attempt to suppress or substitute a thought, a sensory representation or trace of the thought forms. This sensory trace appears to emerge even when individuals report thought suppression success, suggesting the intriguing possibility that non-conscious sensory representations trigger thought control failure.

Our results reveal individuals vary in their ability to control visual thoughts. We found that the average Thought Control Index score was not significantly above zero (*Supplementary Figure 5*), indicating that individuals were, on average, poor at controlling their visual thoughts with thought suppression. Interestingly, individuals with high anxiety and schizotypy showed the lowest levels of thought control, while individuals with high levels of mindfulness exhibited the most thought control (*Figure 3*).

Interestingly, individuals with high anxiety or schizotypy traits showed negative Thought Control Index scores, which implies that the sensory trace of suppressed thoughts was stronger than that of intentionally imagined thoughts. In other words, for these individuals, attempts at thought suppression failed to the degree that the suppressed thought generated a *stronger* sensory trace than an actively imagined thought. This suggests that deficiencies in thought control for highly anxious or schizotypic individuals may arise, in part, from a lack of control over the sensory representations of visual thoughts. On the other hand, individuals with high levels of mindfulness showed positive Thought Control Index scores implying the sensory strength of suppressed thoughts was weaker than that of imagined thoughts reflecting the effectiveness of mindfulness in improving mental control [29].

Our study offers a novel and unique method to objectively measure thought control using binocular rivalry. By probing the sensory basis of emergent thoughts, we provide a simple and inexpensive way to examine the underlying mechanisms of thought control and its failure. In addition, our measure enables examination into the metacognition of thought control, as well as into thoughts which exist in the absence of subjective awareness. We believe this paradigm will be important in objectively measuring thought control, a domain which has thus far been difficult to investigate due to its inherent subjectivity.

Future research should explore the underlying neural mechanism behind thought control and its failure and why such mechanisms go awry in particular populations. Such mechanistic understanding will provide new avenues for clinical treatments. In sum, our study introduces the first objective sensory based method to investigate thought control and provides data to suggest thought control failure may emerge from non-conscious representations in sensory cortices.

## Acknowledgements

This work was supported by Australian NHMRC grants APP1024800, APP1046198 and APP1085404; and a Career Development Fellowship APP1049596 (JP); and an ARC discovery project DP140101560.

## Supplementary Figures

**Supplementary Figure S1.**
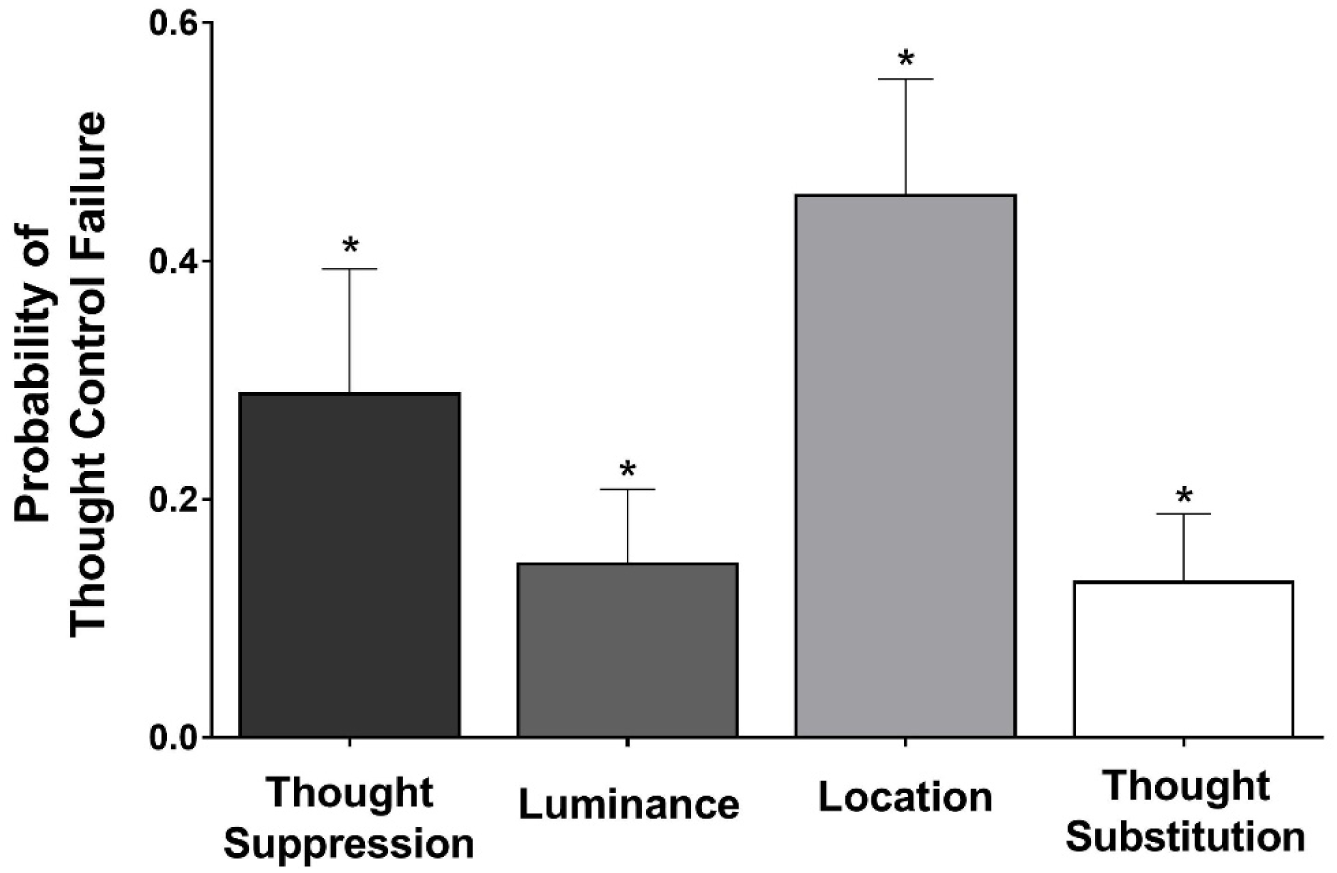
Probability of thought control failure. Thought control failure was measured by a button press during the thought control period in each trial. The probability of thought control failure in all experiments were significantly above zero indicating individuals were unable to successfully control the thought stimuli on a significant number of trials. Error bars represent standard error of the mean (+SEM). Multiple comparisons were corrected by controlling the false discovery rate (FDR) at *q*=0.05.

**Supplementary Figure S2.**
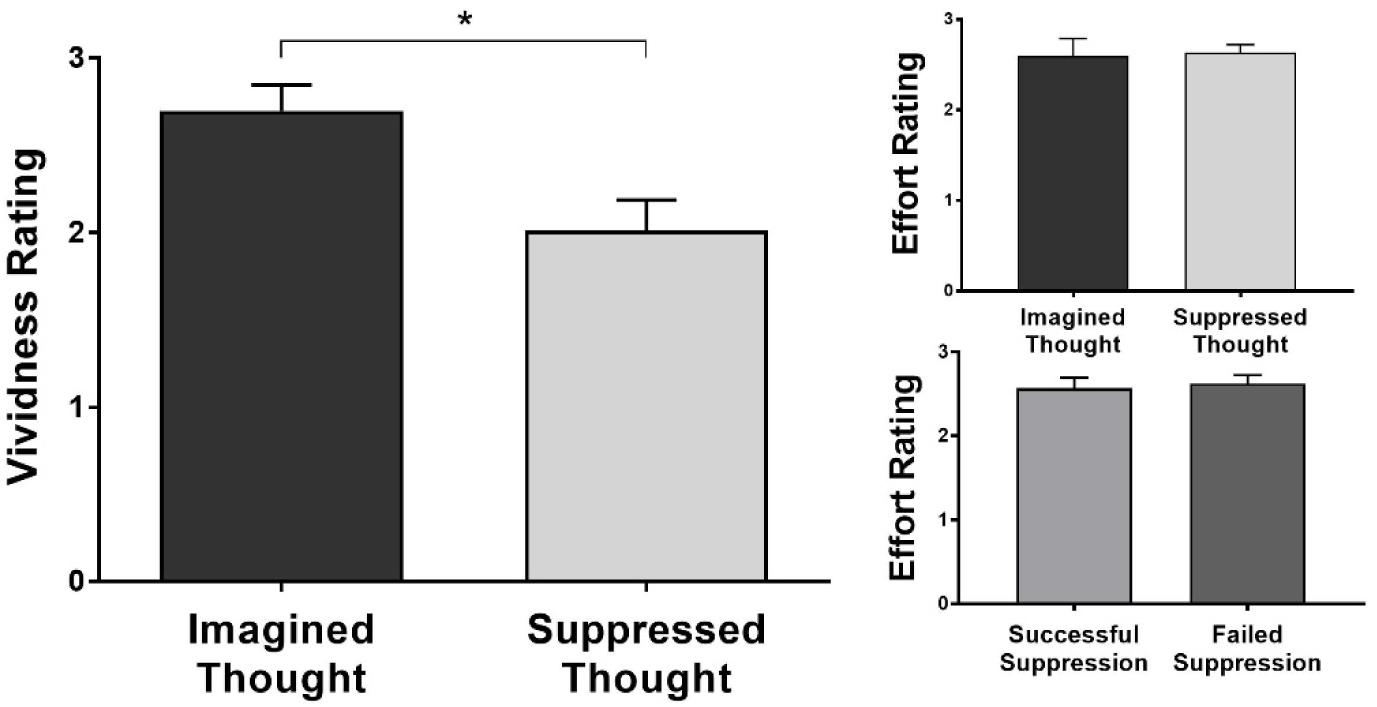
Vividness and effort. In the location experiment, participants rated the vividness and effortfulness of their imagined and suppressed thoughts on a scale of 1-4 (1=not vivid/effortful, 4=very vivid/effortful). Vividness of suppressed thoughts was significantly weaker than imagined thoughts. No difference in effort was found between imagery and suppression trials, and between successful and failed suppression attempts. Error bars represent standard error of the mean (+SEM). Multiple comparisons were corrected by controlling the false discovery rate (FDR) at *q*=0.05.

**Supplementary Figure S3.**
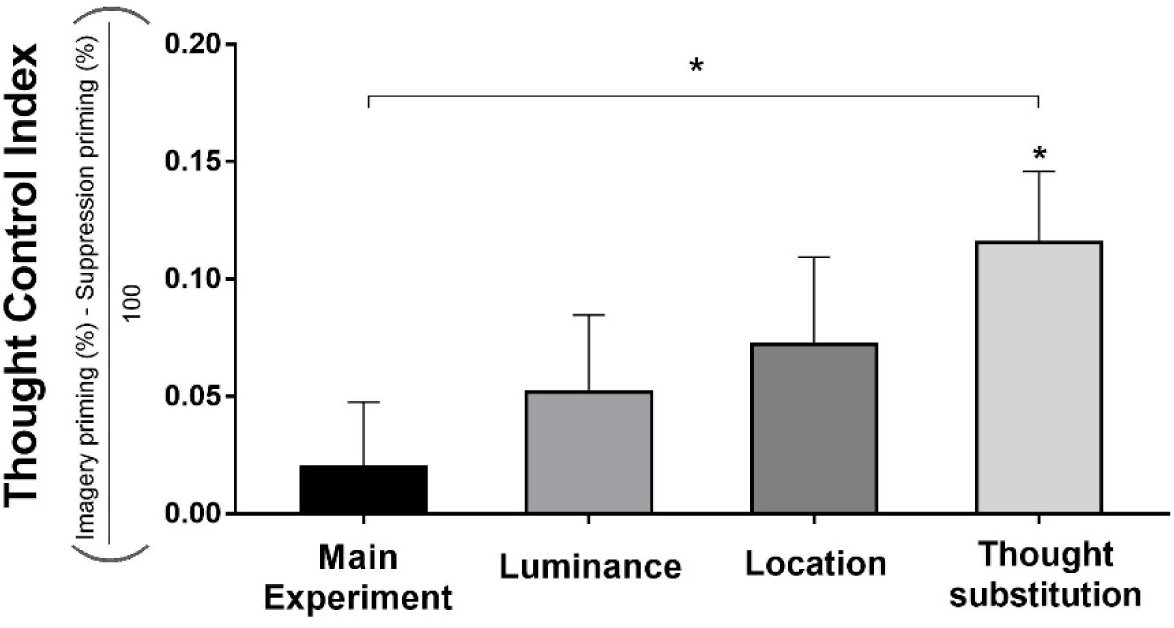
Thought Control Index scores across experiments. A measure of thought control ability was obtained through a devised Thought Control Index. Thought Control Index scores were significantly higher during thought suppression compared to thought suppression and was significantly above zero. Error bars represent standard error of the mean (+SEM). Multiple comparisons were corrected by controlling the false discovery rate (FDR) at *q*=0.05.

**Supplementary Figure S4.**
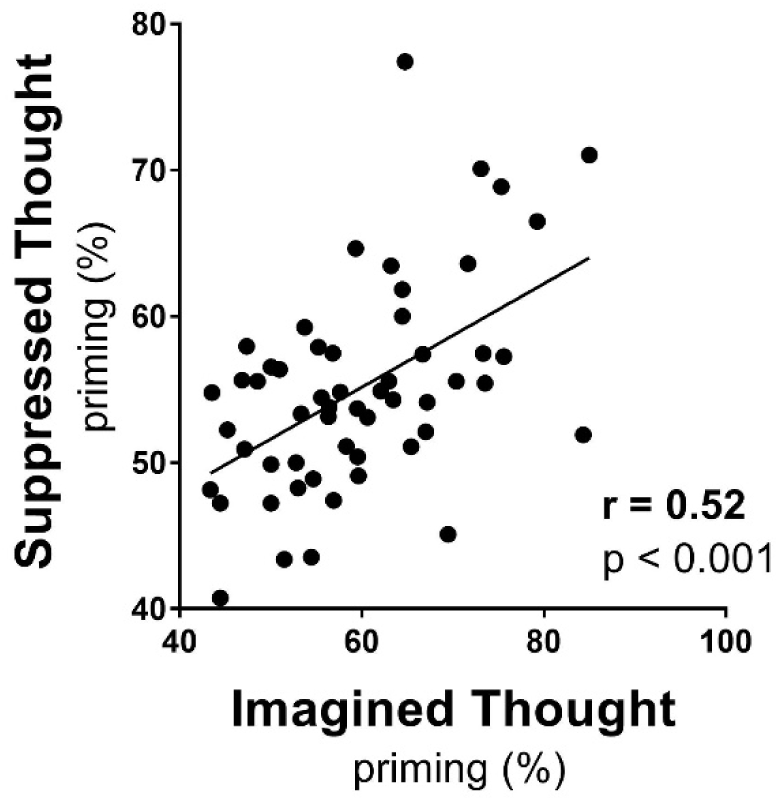
Imagery priming vs suppression priming (n=56). Imagery priming and thought suppression priming showed a significant positive correlation, indicating the sensory strength of suppressed thoughts was linked to mental imagery strength.

**Supplementary Figure S5.**
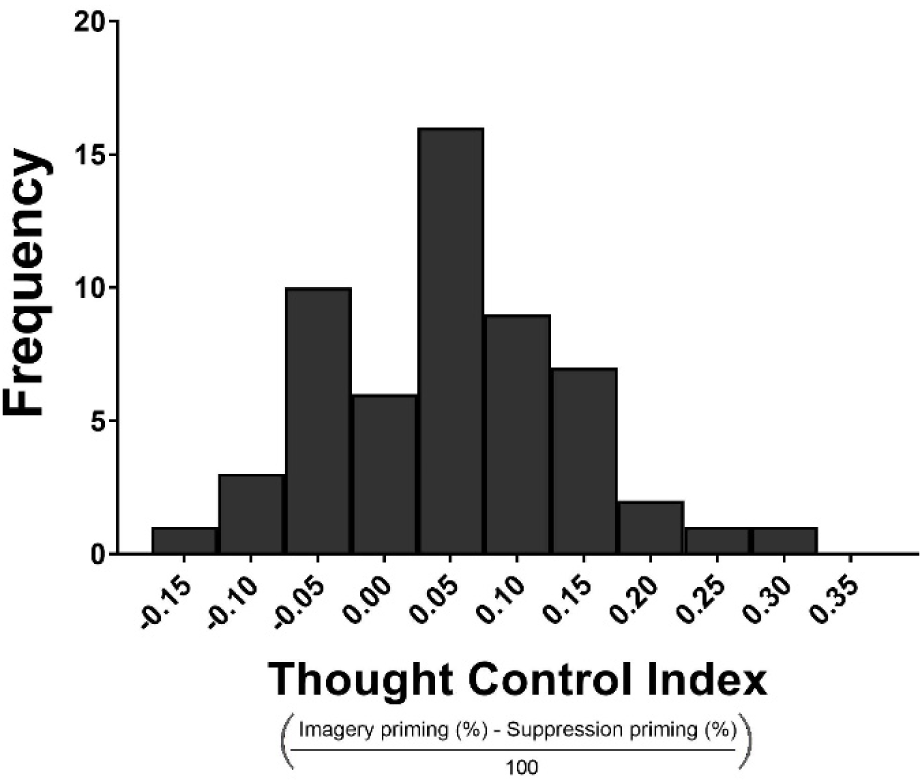
Frequency histogram of Thought Control Index (n=56). Individual scores on the Thought Control Index followed a normal distribution centred around a mean score of 0.045 which was not significantly above zero.

**Supplementary Figure S6.**
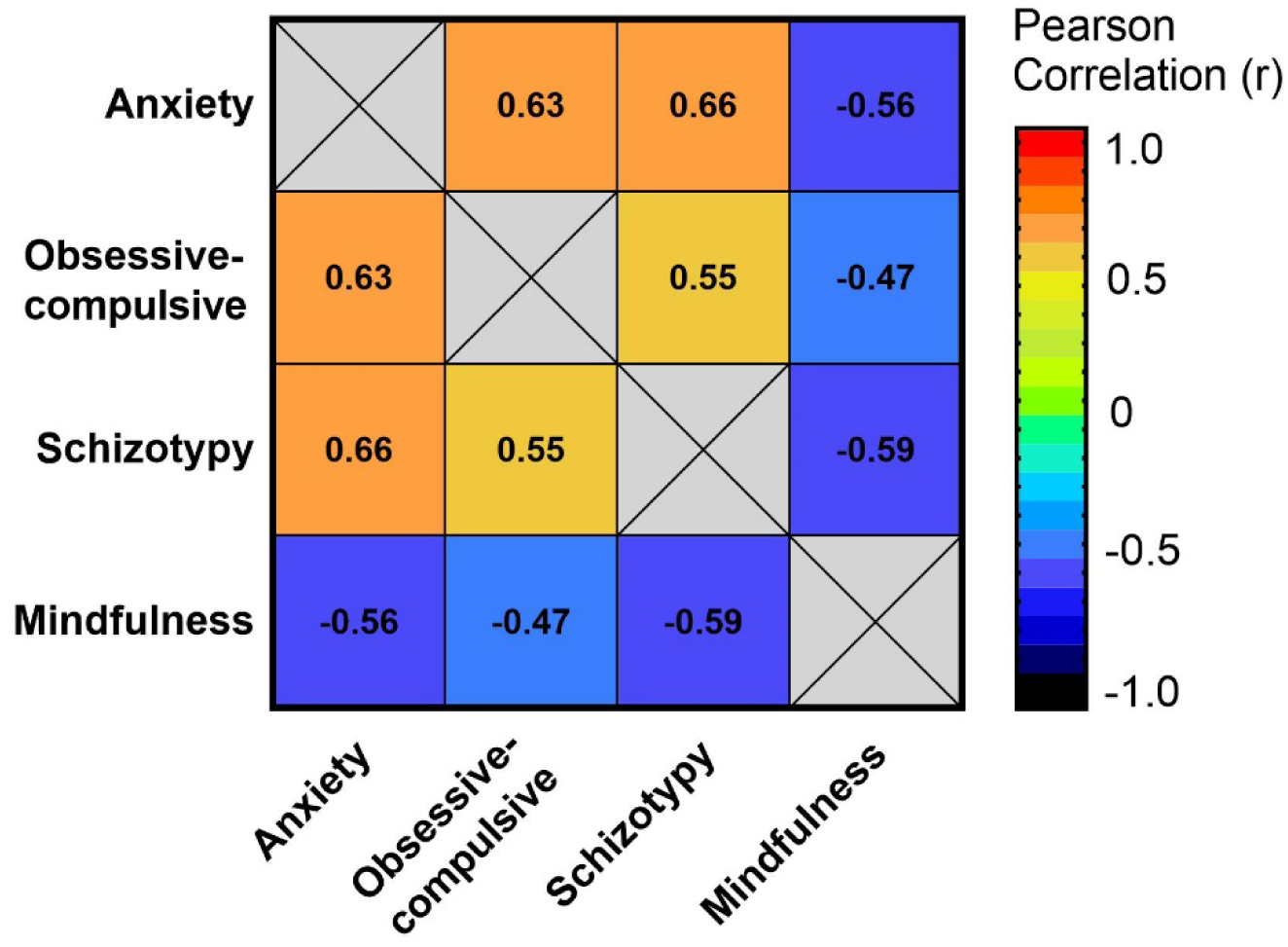
Correlation matrix of questionnaires. Participant scores on each of the four questionnaires were correlated with each other. Numbers inside the boxes denote the Pearson correlation coefficient (r) for the correlation. As expected, significant correlations were found between questionnaire scores. Positive relationships were found between anxiety, obsessive-compulsive tendencies and schizotypy, while mindfulness was inversely correlated with these traits.

